# What do zebrafish prefer? Directional and color preferences in maze tasks

**DOI:** 10.1101/2021.12.22.473814

**Authors:** Matheus Marcon, Radharani Benvenutti, Matheus Gallas-Lopes, Ana Paula Herrmann, Angelo Piato

**Author notes:** Correspondence: Angelo Piato, Ph.D., Programa de Pós-graduação em Neurociências, Instituto de Ciências Básicas da Saúde, Universidade Federal do Rio Grande do Sul (UFRGS), Av. Sarmento Leite, 500/305, Porto Alegre, RS, 90050-170, Brazil. These authors contributed equally to this manuscript (Matheus Marcon, Radharani Benvenutti).

## Abstract

Studies regarding the animals’ innate preferences help elucidate and avoid probable sources of bias and serve as a reference to improve and develop new behavioral tasks. In zebrafish research, the results of innate directional and color preferences are often not replicated between research groups or even inside the same laboratory raising huge concerns on the replicability and reproducibility. Thus, this study aimed to investigate the male and female zebrafish innate directional and color preferences in the plus-maze and T-maze behavioral tasks. As revealed by the percentage of time spent in each zone of the maze, our results showed that males and females zebrafish demonstrated no difference in directional preference in the plus-maze task. Surprisingly, male and female zebrafish showed color preference differences in the plus-maze task; males did not show any color preference, while female zebrafish demonstrated a red preference compared to white, blue, and yellow colors. Moreover, both male and female zebrafish demonstrated a strong black color preference compared to the white color in the T-maze task. Thus, our results demonstrate the importance of innate preference assays involved with the directionality of the apparatus or the application of colors as a screening process conducting behavioral tests (e.g., anxiety, learning and memory assessment, locomotion, and preference) and highlight the need to analyze sex differences.

## 1 INTRODUCTION

Behavioral neuroscience research is fundamental and provides essential findings to help understand and interpret human behavioral phenotypes. Also, it provides researchers knowledge regarding the neural bases of behaviors, including those subjacent to neuropsychiatric disorders and drug effects on the central nervous system [1–3]. In pre-clinical research, experimental models have been used to develop, model, and monitor diseases’ progress, favoring progress in understanding the neurobiology of human diseases [4,5]. Many behavioral tests are employed to assess animal behavior. They range from tests that evaluate less complex behaviors (locomotor assessment) to those that evaluate more complex behaviors (learning and memory assessment) [6–8].

Mazes are experimental tools often implemented in behavioral tests, once they are applicable across species and, with small changes to its configuration, allow to evaluate different sets of behavioral paradigms (e.g., anxiety, learning and memory assessment, locomotion, and preference) [9–13]. Most of the data generated using these apparatus come from studies with rodent models, generally seeking to assess anxiety or cognition [13,14].

Along with the vast number of tasks designed for rodents, researchers also explored the versatility of the mazes adapting these apparatus to assess behavioral data from many other model organisms such as fruit flies [15], frogs [16], and fish [17]. For example, in zebrafish, mazes have been used to study anxiety [12,18,19], learning and memory [20–22], locomotion [23–25], and preference [26–29]. Recently, a review article featuring an overview of maze apparatuses and protocols to assess zebrafish behavior was published by our research group and now is available in the literature [30].

Zebrafish is a model organism increasingly being used in behavioral neuroscience research, enabling the study of a vast range of behavioral paradigms [5,31], such as anxiety [32], learning and memory [33], and seizure [34]. Furthermore, the zebrafish is a successful model for translational research on human neurological disorders [35] and high-throughput screening of potential treatments [36]. These animals provide rational, quick, and low-cost tools to research due to their genetic tractability, conserved neurobiology, as well as evident behavioral responses essential to model neuropsychiatric-like disease phenotypes [37].

The zebrafish innate directional and color preferences also are frequent subjects of scientific scrutiny [26,38–41], but results are often not replicated between laboratories or even in the same research group [42]. For example, results show several inconsistencies in the color preference studies regarding the fish preference or aversion by the same color [38,39,43]. It can be related to methodological problems such as the lack of standardized protocols, raising huge concerns on several studies’ replicability and reproducibility [44].

Studies about the animals’ innate preference not only help to elucidate and avoid probable sources of bias (e.g., zebrafish directional preference can be the reason for the fish to spend more time in one of the arms of the maze blunting the analysis of any intervention), but also serve as a reference to improve and/or develop new behavioral tasks. Learning and memory protocols, for example, often implement the technique of pairing rewards stimulus (e.g., food or conspecifics) with colorful visual cues [39,45], while anxiety protocols mostly using the black and white colors to determine anxiety-like phenotypes based on scototaxis [46–48], pointing once again to the importance of detecting zebrafish preference or avoidance for different colors. Several behavioral studies showed male and female differences for aggressiveness [49], stress [50], and drug response [51], highlighting the relevance to consider all behavior analyzes of manner sex-dependent.

In this context, this study aimed to investigate the male and female zebrafish innate directional and color preferences in the T-maze and plus-maze behavioral tasks to identify possible sources of bias and provide insights that may contribute to the standardization of future protocols.

## 2 MATERIAL AND METHODS

### 2.1 Animals

All experiments were performed using 60 adult short-fin wild-type zebrafish (*Danio rerio*, 6-month-old, 3–4 cm long, weighing 400-500 g) in a 50:50 male/female ratio. The fish were obtained from a local commercial supplier (Delphis, RS, Brazil) and maintained for at least 15 days in an animal facility (Altamar, SP, Brazil) before being assigned to the experimental tanks. The density of animals was maintained at a maximum of 2 animals per L. The water of the recirculation system was kept in the conditions required for the species (27 ± 1°C; dissolved oxygen at 7.0 ± 0.4 mg/L; pH 7.0 ± 0.3; total ammonia at <0.01 mg/L; alkalinity at 22 mg/L CaCO_3_; total hardness at 5.8 mg/L; and conductivity of 1500–1600 μS/cm) being constantly filtered by mechanical, biological and chemical filters. Animals were fed twice a day (09:00 a.m./05:00 p.m.) with commercial flake food (Poytara^®^, Brazil) plus the brine shrimp *Artemia salina*. All tests performed in this study followed ARRIVE guidelines [52]. Lighting conditions consisted of a light/dark cycle of 14/10 hours. At the end of the experiments, zebrafish were euthanized by immersion in cold water (0 to 4 °C) until cessation of the any movements, followed by decapitation to ensure death according to the AVMA Guidelines for the Euthanasia of Animals [53]. All procedures were approved by the institutional animal welfare and ethical review committee (approval n° 36248/2019).

### 2.2 Experiment design

This study consisted of 3 independent experiments with maze tasks. All our results were replicated and confirmed by two independent experiments for each of the 3 experiments. In each experiment, one different set of animals was used after the 15 days of the acclimation period to laboratory conditions. One single experimental group (n=20) was allocated in two independent experimental tanks (A and B) of 16-L (40 × 20 × 24 cm) where stayed for 7 days before the start of the experiments and throughout the experimental period. In the experimental tanks, the animals were fed twice a day (in the morning after the experiments and at 05:00 p.m.). Block randomization procedures were used to counterbalance the sex of the animals and the two independent experimental tanks, the order of the animals tested, and the maze and color positions during the tests. The sample size used in the present study was defined a priori based on previous literature and pilot studies. 20 fish (10 males and 10 females) were used in each experiment. One exclusion criterion was established prior, in which subjects that frequently stopped (more than 50% of the test time) or never swam would be excluded from the data analysis. Thus, at the end of all experiments, the number of animals was not the same for all experiments.

Specifically, in experiment 1, the number of animals was reduced to two males (one male died during the habituation phase, and one male was excluded from the data analysis) and two females (excluded from the data analysis). In experiment 2, the number of animals was bigger for females than for male sex (at the end of all experiments, when sex was confirmed by dissection, we observed more females than males in the experimental tanks). In experiment 3, the number of animals was reduced to one male (one male died on the test day) and one female (excluded from the data analysis). The tanks were filled with water from the animal facility. The determination of the sex of the animals was performed by dissection, followed by the analysis of the gonads.

### 2.3 Maze design

We have used the same apparatus for all experiments. The complete maze design utilized in this study is represented in figure 1. The apparatus consisted of transparent plexiglass (1 cm thick) cross-shaped maze with a start zone (10 × 10 × 15 cm) into the stem arm (40 × 10 × 15 cm) and 3 short arms (20 × 10 × 15 cm) connected to the final stem arm. This apparatus is easily adaptable to different maze shapes such as plus or T-mazes (implemented in these experiments) by closing sliding doors present along the entire maze every 10 cm. The apparatus was placed inside a white plastic box (93 × 55 × 58) that contained support of white plexiglass attached to the two sides of the box and served to suspend a camera on top of the apparatus, which allowed filming the behavior during the test from above. The camera distance from the floor of the box was 89 cm. The box was covered with a white fabric to avoid interference by environmental cues. A source of light (LED strip light) was fixed 5 cm above the box floor around all inside walls to ensure that lighting conditions were the same in each of the arms apparatus (275 lux was measured across the entire maze with the aid of the Lux Light Meter Pro application version 2.0). The water level was set at 5 cm inside the maze, and the water temperature was maintained at 27°C (± 1°C) throughout the entire experiment. The heater wire was covered with white adhesive tape (same color as the box and white fabric) to avoid environmental clues. The maze was emptied and cleaned between the test of each animal.

**Figure 1:**
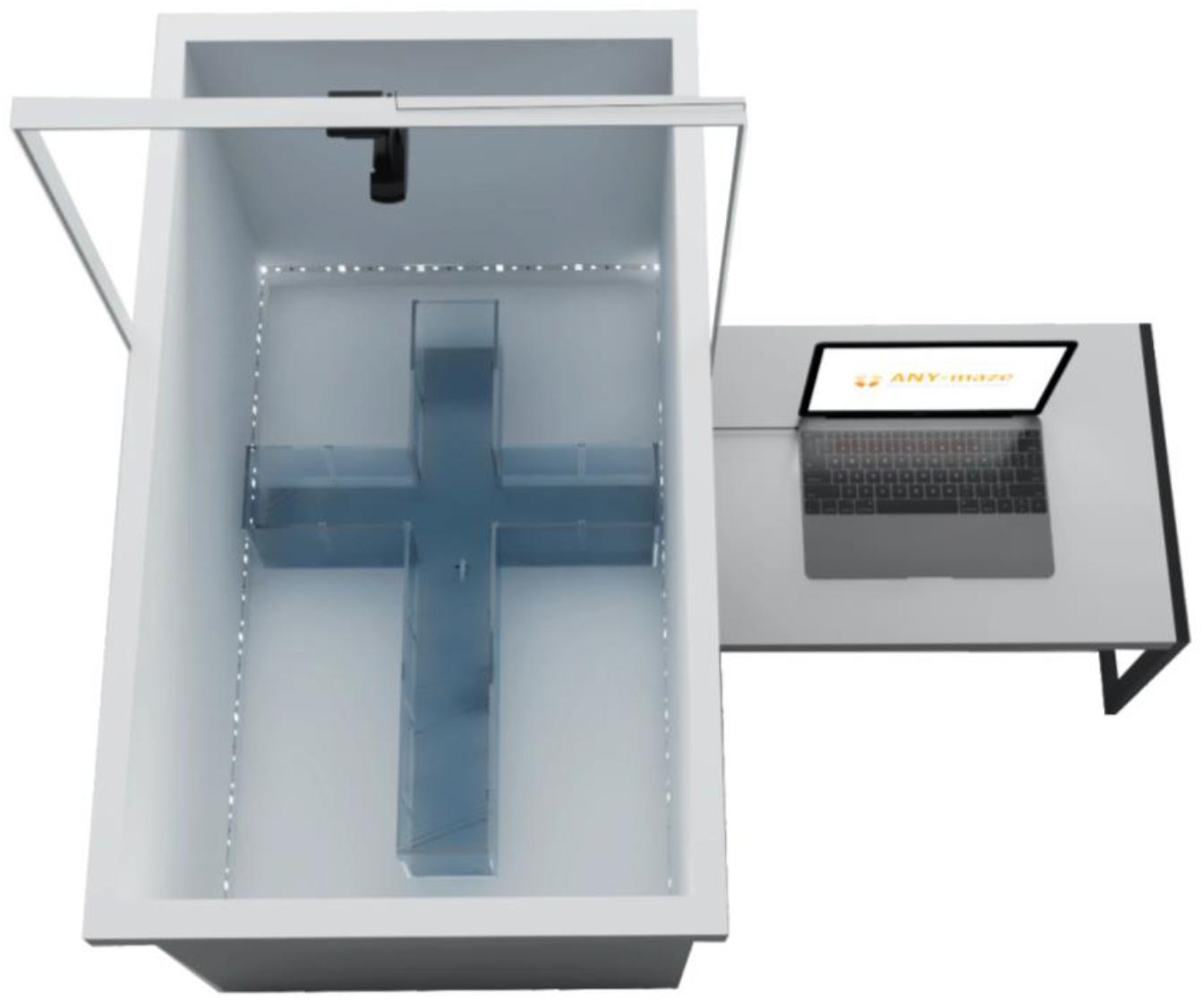
3D Illustrative representation of the maze apparatus and the experimental configuration used in behavior tests performed in this study.

### 2.4 Habituation and task protocol

All tasks were performed between 08:00 and 12:00 a.m. in a room used exclusively for experiments with mazes, which was a different room than the one where the animals were housed. To avoid novelty stress induced by the environment, all animals were transported to the behavior room 1 hour before the tests. The maze tasks consisted of 5 days. In the first 4 days, the fish were placed in the apparatus in groups for habituation. The number of animals gradually decreased over the days, helping to minimize social and novelty stress (Sison and Gerlai 2010). On the fifth day (last day), the fish were tested individually in the maze task. Briefly, on the first day of habituation, all animals of the same group were placed in the apparatus’s start zone. For the plus-maze task, the start zone was the center zone, and for the T-maze task, the start zone was at the beginning of the stem arm. After the start zone doors were opened, the fish freely explored the apparatus for 20 minutes. On the second day, the number of animals was reduced by half, and the fish could explore the maze for 10 min. By the third day of habituation, the number of animals was again reduced by half, and fish could swim for only 5 min. On the fourth day (last day for habituation), fish were individually placed on the maze and explored freely for 5 min. On the fifth day (test day), the fish were individually placed on the apparatus’s starting zone remaining there for 2 min to habituate. Posteriorly, the doors of the starting zone were opened the fish explored the maze for 5 min.

### 2.5 Behavioral analyses

On the fifth day (behavior assessment), the animals were not fed. Following a protocol previously elaborated with randomization procedures using random.org software (computerized random numbers) to avoid potential confounders, the animals were transported from the experimental tank (A or B) to the test. Animal behavior was recorded with a webcam (Logitech^®^ C920 HD pro) from above. The behavior analyses were performed from the recorded videos dividing the tank into virtual zones with ANY-Maze^®^ automated tracking software 4.99 version (Stoelting Co., Wood Dale, IL, USA) for Windows system 10 version. For the behavioral and statistical analyses, blinding was achieved by assigning to each animal a code that was revealed only after data analyses (the coding was performed by a researcher who did not participate in the experiments).

### 2.6 Directional preference

In experiment 1, to assess directional preference, our maze has been configured to take the shape of a plus-maze. For this, the labyrinth stem arm (40 cm) was blocked by closing a sliding door at half the length of the arm.

Thus, the maze stayed with four equal arms (each 20 cm long). We positioned the maze with the help of a compass (available on the iOS version 13.5.1) so that each arm was pointed in one of the cardinal directions (north, south, east, and west). Briefly, the features of the plus-maze consisted of one center zone (10 × 10 × 15 cm) and four identical arms (20 × 10 × 15 cm). The maze position was counterbalanced (turning the maze by 90°) between animals to avoid possible biases. Afterward, behavioral analyses were performed based on the recorded videos’ analysis by virtually dividing the maze into four zones (north zone, south zone, east zone, and west zone). The time spent in each zone was used as the exploratory parameter and expressed in percentage (%).

### 2.7 Color preference

In experiment 2, the same plus-maze used for experiment 1 was implemented to assess zebrafish’ innate color preference. For this purpose, each arm was covered with a colored sleeve (white, red, blue, or yellow). The position of each sleeve was counterbalanced between animals to avoid possible biases. Afterward, behavioral analyses were performed based on the recorded videos’ analysis by virtually dividing the maze into four zones (white zone, red zone, blue zone, and yellow zone). The time spent in each zone was used as the exploratory parameter and expressed in percentage (%).

### 2.8 Black or white color preference

In experiment 3, to assess innate zebrafish preference between the color black or white, our maze has been configured to take the shape of a T-maze. One of the three short arms of the maze was blocked by closing a sliding door. Thus, the maze stayed with two short arms (20 cm length) and the stem arm (40 cm length). We covered one of the short arms with a black color sleeve and the other with a white color sleeve. The position of each sleeve was counterbalanced between animals. Briefly, the maze consisted of one start zone (10 × 10 × 15 cm) into the stem arm (40 × 10 × 15 cm) and two identical short arms (20 × 10 × 15 cm) connected to the final stem arm. Afterward, behavioral analyses were performed out based on the recorded videos’ analysis by virtually dividing the maze into four zones (start arm, neutral zone, white zone, and black zone). The time spent in each zone was used as the exploratory parameter and expressed in percentage (%).

### 2.9 Statistical analysis

Results were analyzed by generalized estimating equation (GEE) followed by Bonferroni post hoc test when appropriate. Subjects from the same experimental group but different experimental tanks did not differ in any behavioral measures, so they were combined into a male and female group for statistical analysis and results presentation. The evaluation of the data distribution for each variable was performed through the residual analysis.

When the normal distribution was not adequate, other distributions/transformations were considered (Gamma and Log-normal distribution). The differences were considered significant at p<0.05. The data were expressed as the mean ± standard error of the mean (S.E.M). Data were analyzed using IBM SPSS Statistics 18.0 for Windows 10 version, and the graphics were assembled with the GraphPad Prism version 8 for macOS Big Sur 11.0.1 version.

## 3 RESULTS

Figure 2 shows the zebrafish innate directional preference in the plus-maze task positioned with each arm pointing to one of the cardinal directions (north, south, east, and west). The GEE found no significant interaction between sex and direction (χ^2^=3.467; 3; p=0.325). There were no statistical differences in time spent by the male zebrafish (χ^2^=6.419; 3; p=0.093) and female zebrafish (χ^2^=2.282; 3; p=0.516) between each of the zones. Therefore, both sexes of zebrafish did not show a directional preference in this task.

**Figure 2.**
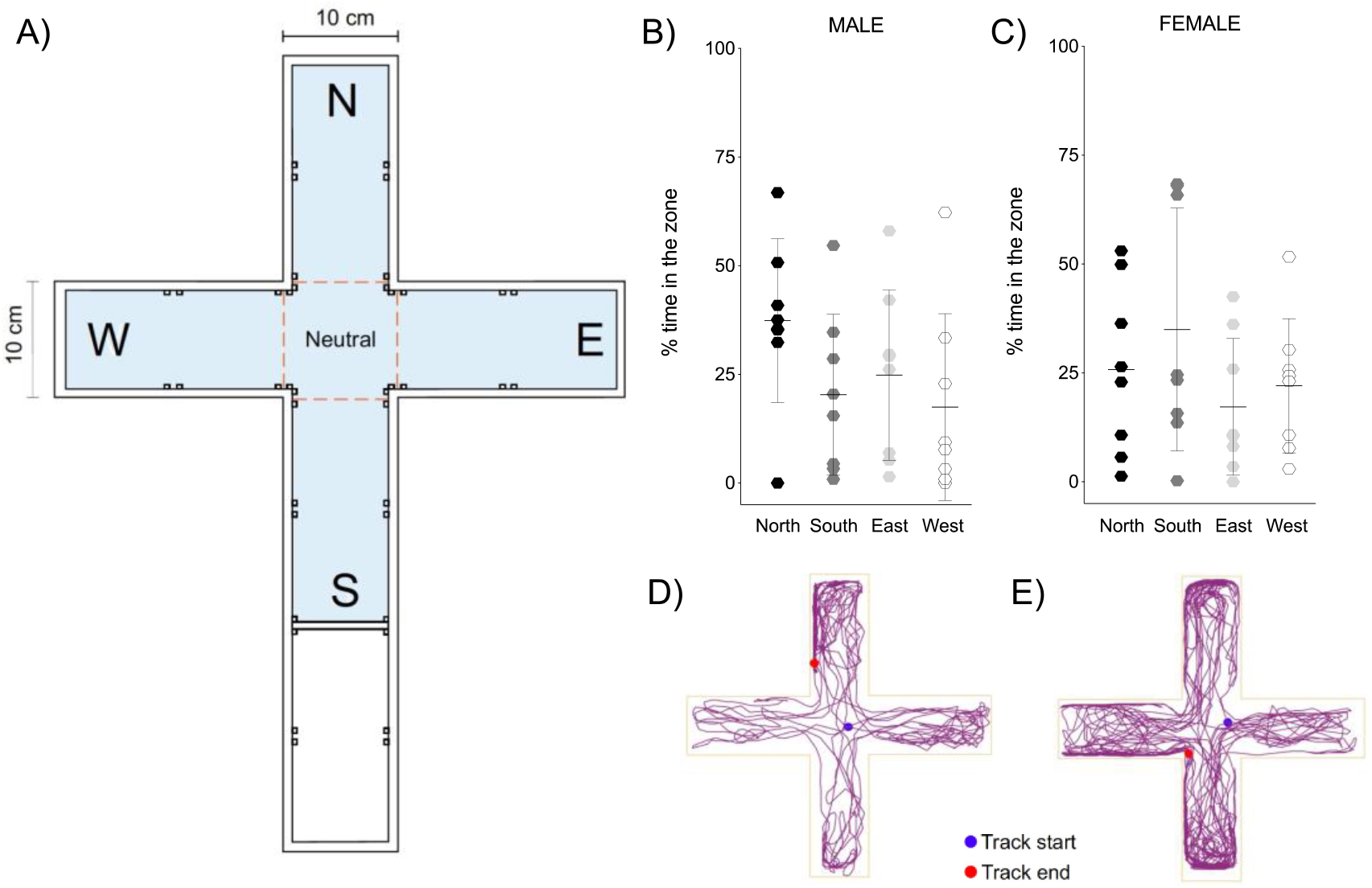
The zebrafish innate directional preference in the plus-maze task was positioned with each arm pointing to one of the cardinal directions (north, south, east, and west). (A) Representative plus-maze design; (B) Male and (C) female % of time spent in each zone of the maze; (D) Male and (E) female representative track plot of the one animal behavior from the group for 5 min. Male n = 8; Female n = 8. Data are expressed as a mean ± S.E.M. Generalized estimating equation (GEE).

Figure 3 shows the zebrafish innate color preference in the plus-maze task with each arm of the maze covered with a colored sleeve (white, red, blue, or yellow). The GEE revealed an interaction between sex and colors (χ^2^=9.774; 3; p=0.021). Thus, male and female zebrafish showed differences in preferences for primary colors. There were no statistical differences (χ^2^=7.203; 3; p=0.066) in the time spent in each of the zones in the male zebrafish. However, it was revealed that the female zebrafish spend more time in the red zone than the white, blue, and yellow zones (χ^2^=18.730; 3; p<0.001).

**Figure 3:**
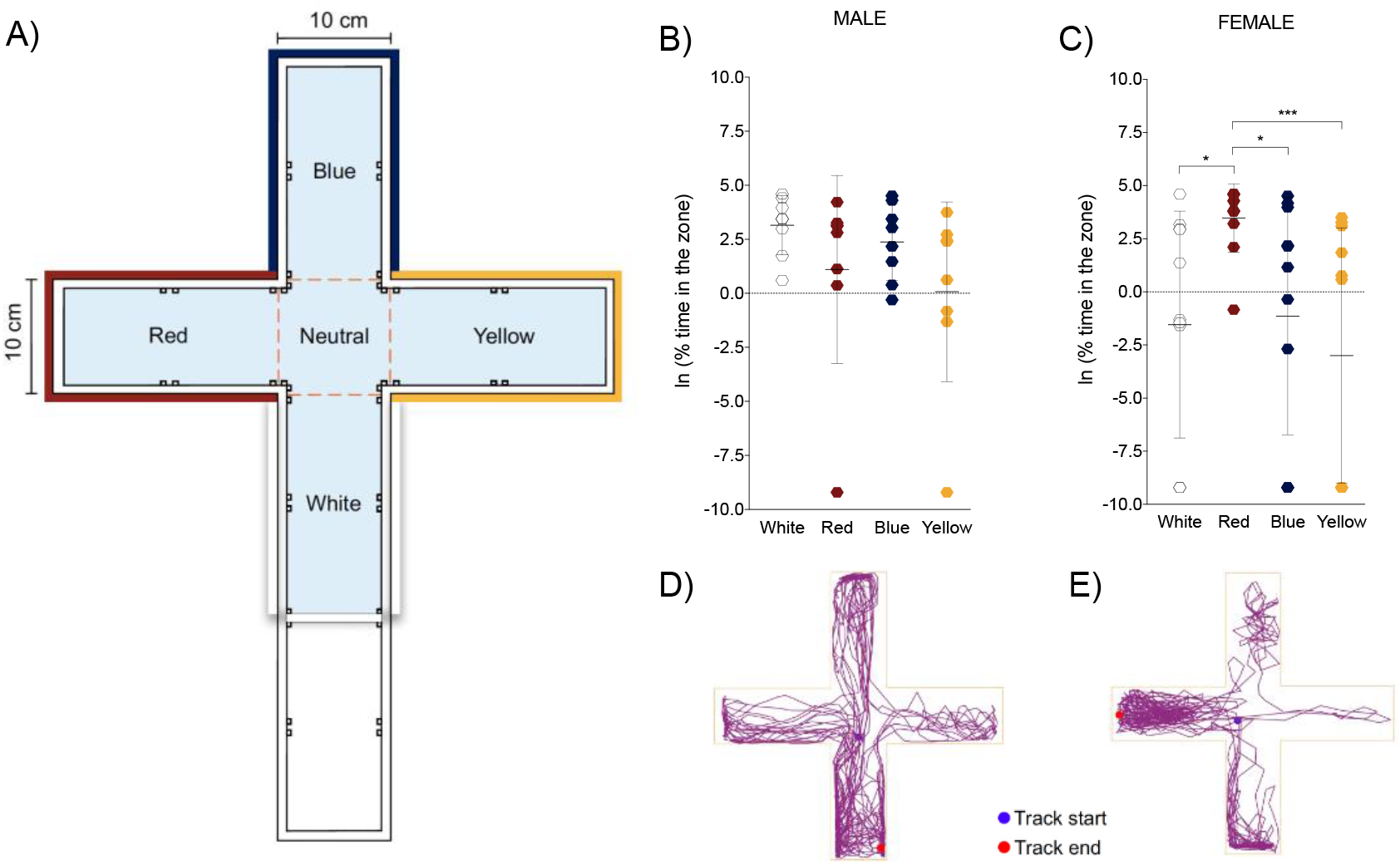
The zebrafish innate color preference in the plus-maze task with each arm of the maze covered with a colored sleeve (white, red, blue, or yellow). (A) Representative plus-maze design; (B) Male and (C) Female % of time spent in each zone of the maze; (D) Male and (E) Female representative track plot of the one animal behavior from the group for 5 min. Male n = 8; Female n = 11. Data are expressed as a mean ± S.E.M. Generalized estimating equation (GEE) followed by Bonferroni post hoc test. *p<0.05; ***p<0.001.

Figure 4 shows the innate zebrafish preference between the color black or white in the T-maze task with each short arm of the maze covered with sleeves of the black or white color. The GEE revealed no significant interaction between sex and colors (χ^2^=1.745; 2; p=0.418). Male (χ^2^=896.319; 2; p<0.0001) and female (χ^2^=462.796; 2; p<0.0001) zebrafish showed a strong preference for the black color when compared to the white color. Therefore, both sexes spent more time in the black zone when compared to the white zone.

**Figure 4:**
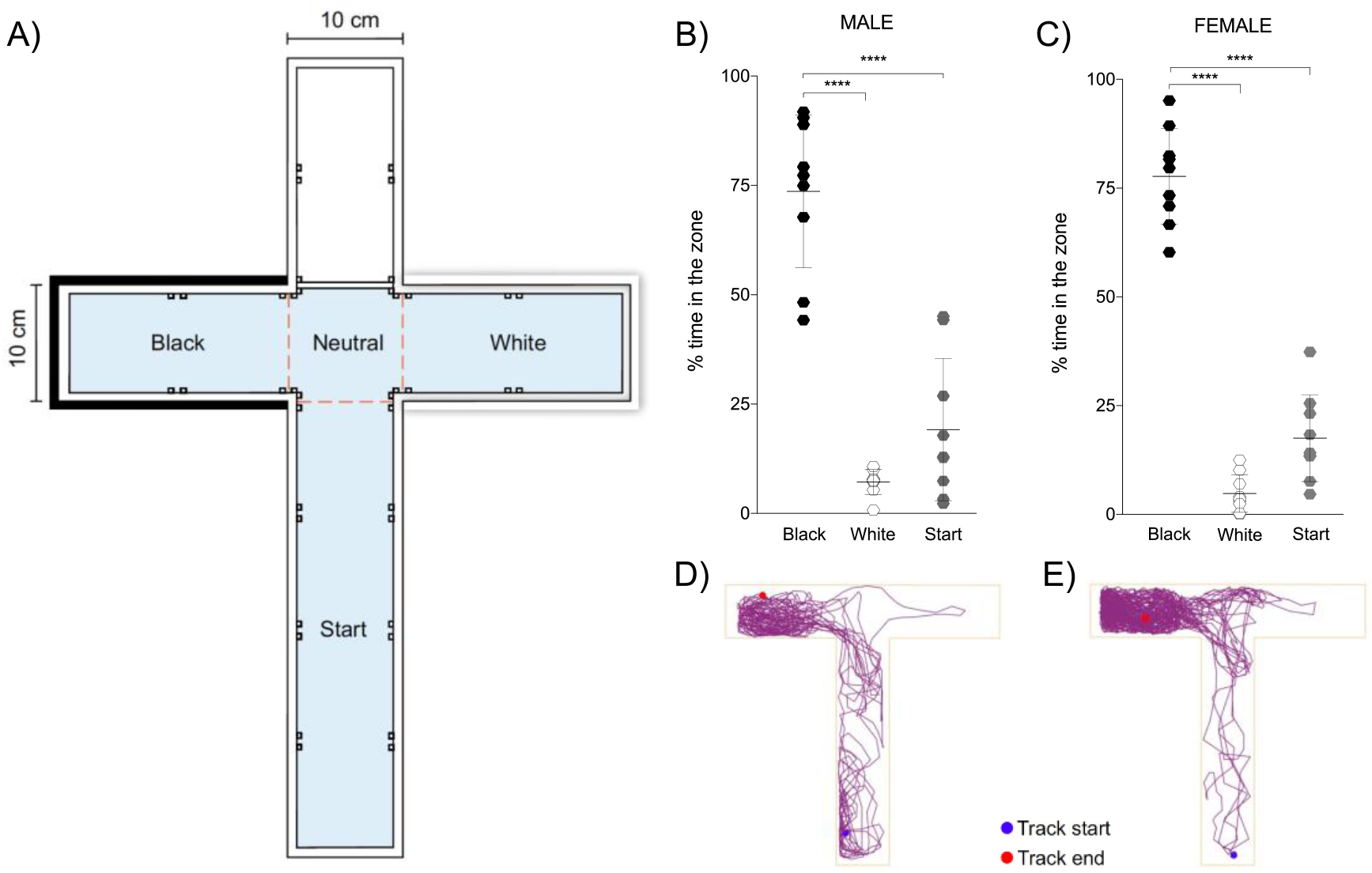
The innate zebrafish preference between the color black or white in the T-maze task with each short arm of the maze covered with the black or white color sleeves. (A) Representative T-maze design; (B) Male and (C) Female % of time spent in each zone of the maze; (D) Male and (E) Female Representative track plot of the behavior of one animal from the group for 5 min. Male n = 9. Female n = 9. Data are expressed as a mean ± S.E.M. Generalized estimating equation (GEE) followed by Bonferroni post hoc test. ****p<0.0001.

## 4 DISCUSSION

In this study, we have investigated the zebrafish innate directional and color preferences in the maze’s tasks to identify possible biases and provide results that contribute to the standardization of future protocols. Our results revealed that male and female zebrafish had no directional preference, and both sexes showed a similar preference in the plus-maze task. Still, male and female zebrafish showed color preference differences in the plus-maze task; males did not show any color preference, while females preferred the red color compared to the white color, blue color, and yellow color. Moreover, male and female zebrafish showed no differences in black and white color preference; both sexes showed a preference for the black color when compared to the white color in the T-maze task.

### 4.1 Directional preference

The analysis of directional preference contributes as a screening process when it is desired to carry out behavioral tests (for example, other preferences or learning and memory), mainly in maze tasks. In experiment 1, the plus-maze task, with each arm of the maze pointing to the cardinal directions (north, south, east, and west), was used to assess the zebrafish’s directional preference. The plus-maze task is ideal for assessing directional preference because the maze with four equal arms allows researchers to point the apparatus to each of the main directions.

Our result showed that zebrafish of both sexes had no directional preference; both sexes similarly had directional behavior, as demonstrated by the % of time spent in each zone of the maze. Our result is different from another literature study that showed bimodal directional preference (east-west) when the plus-maze was positioned pointed to the same directions (cardinal points) [40]. However, agreeing with the authors of this study’s conclusion, we hypothesized that these differences could be explained by the differences in the protocols (for example, habituation to maze), mainly related to the labyrinth’s dimensions. In our study, the dimension of plus-maze was the same type used in the behavior analysis (20 × 10 cm) (see for example [18,45], while in the study of Osipova et al., (2016) the plus-maze dimension was smaller (6 × 3 cm).

When the zebrafish directional preference was tested in the T-maze task with the two short arms pointing northeast/southwest or north/south, males zebrafish showed a directional preference to southwest and south directions, respectively, but females had no directional preference [54]. Therefore, biological differences between the sexes can contribute as a relevant factor in the behavioral analysis, but this was not observed in our results [55]. Although the T-maze task is not ideal for analyzing directional preference, it can be useful as screening before other behavior tests in this maze to avoid direction bias.

### 4.2 Color preference

Several behavioral protocols that assess zebrafish learning and memory use colored clues as a conditioned stimulus. Despite this, there is no consensus in the scientific literature regarding the zebrafish’ innate color preference. For example, two studies found that zebrafish showed a greater preference for red over yellow [38,39], while another study found that zebrafish had a preference for blue and green and avoided yellow and red [56,57]. A lack of standardization in the protocols used to assess color preference could explain why there is inconsistency in the scientific literature results. Furthermore, most studies did not evaluate differences between males and females.

For this reason, in experiment 2, we investigated if the zebrafish has an innate color preference, aiming to add relevant information for this research field. For this reason, in experiment 2, we investigated if the zebrafish has an innate color preference, aiming to add relevant information for this research field. Surprisingly, male and female zebrafish showed color preference differences; even though we did not observe innate color preference in males, the females showed a preference for the red color compared to the white color, blue color, and yellow color, as demonstrated by % of time spent in each zone of the maze. It is already known that there are behavioral differences between males and females in terms of aggressiveness [49], stress [50], and drug response [51]. Our study shows a sex difference in innate color preference for the first time, emphasizing the importance of assessing differences between males and females in studies that use colors as clues.

### 4.3 Black or white color preference

In experiment 3, the T-maze task with each short arm of the maze covered with sleeves of the black or white color was used to investigate if the zebrafish has an innate black or white color preference. It is important to differentiate the task implemented in this study from other protocols of light vs. dark preference of zebrafish once some researchers utilize “light” as interchangeable with “white” and “dark” as interchangeable with “black” when these represent two different variables (color of the walls vs. level of illumination of the apparatus) [46]. Another key factor of this test is the habituation period, which reduces the animals’ anxiety as we exclude the novelty factor. In this context, our test focus on zebrafish’ preference rather than anxiety-like behaviors assessed in similar protocols using these colors [32,47,58–60].

Our data showed that male and female zebrafish had the same preference when it was compared between black and white colors; both sexes had a strong preference for the black color over the white color, as shown by % of time spent in each zone of the maze, which replicated findings shown in other studies. The first reports on the strong zebrafish preference for the black color chamber were described by Serra *et al*. (1999), whose results were also replicated by several researchers [46,61], leading to the development of protocols to assess anxiety-like behaviors based on the animals’ scototaxis [47,62].

On the other hand, juvenile zebrafish display a strong avoidance of the black color chamber when facing the same task, possibly due to endogenous avoidance for the dark color at that stage of life [63]. Other researchers also put the black or white paradigm to the test, pointing out some inconsistencies in the methodology implemented, like the background shades, illumination in the testing facility, the settings of the apparatus, and other interferents that may lead researchers to improper interpretation of the results [46,64,65] By manipulating the light level, for example, researchers reported that under different light conditions, zebrafish exhibits a preference for different chambers, either the white or the dark one [66].

Although most of the studies point to a preference for the black color chamber by the zebrafish, it was important to characterize the normal behavior of the zebrafish in the black and white color preference test under the experimental conditions and the maze implemented in our laboratory, improving, by these means, the execution of future experiments through standardization and the avoidance of biases that may interfere in the obtention of behavioral data (e.g., implementing zebrafish models of anxiety).

## 5 CONCLUSIONS

Overall, we have shown in this study that zebrafish had some innate preferences. Male and female zebrafish showed no directional preference in the plus-maze task. However, male and female zebrafish showed a different color preference in the plus-maze task; male zebrafish did not show any color preference, while female zebrafish preferred the red color compared to the white color, blue color, and yellow color. Both sexes showed a strong preference for the black color when compared to the white color in the T-maze task. Our results show the importance of innate preference analysis involved with the directionality of the apparatus or the application of colors as a screening process conducting behavioral tests (e.g., anxiety, learning and memory assessment, locomotion, and preference) and highlight the need to analyze differences between the sexes. This study was confirmatory to characterize the innate directional and color preference of zebrafish, identifying possible biases, and providing insights that contribute to the standardization of future protocols.

## Acknowledgments

We thank the Conselho Nacional de Desenvolvimento Científico e Tecnológico (CNPq, proc. 303343/2020-6), Coordenação de Aperfeiçoamento de Pessoal de Nível Superior - Brasil (CAPES), and Pró-Reitoria de Pesquisa (PROPESQ) at Universidade Federal do Rio Grande do Sul (UFRGS) for funding.

## REFERENCES

[1] E.J. Nestler, S.E. Hyman, Animal models of neuropsychiatric disorders, Nat. Neurosci. 13 (2010) 1161–1169. https://doi.org/10.1038/nn.2647.

[2] C. Maximino, R.X. do C. Silva, S. de N.S. da Silva, L. do S. dos S. Rodrigues, H. Barbosa, T.S. de Carvalho, L.K. dos R. Leão, M.G. Lima, K.R.M. Oliveira, A.M. Herculano, Non-mammalian models in behavioral neuroscience: consequences for biological psychiatry, Front. Behav. Neurosci. 9 (2015) 233. https://doi.org/10.3389/fnbeh.2015.00233.

[3] C. Maximino, F.J. van der Staay, Behavioral models in psychopathology: epistemic and semantic considerations, Behav. Brain Funct. 15 (2019) 1. https://doi.org/10.1186/s12993-019-0152-4.

[4] P.M. Conn, Animal models for the study of human disease, 2nd ed., Academic Press, 2017.

[5] A.M. Stewart, O. Braubach, J. Spitsbergen, R. Gerlai, A.V. Kalueff, Zebrafish models for translational neuroscience research: from tank to bedside, Trends Neurosci. 37 (2014) 264–278. https://doi.org/10.1016/j.tins.2014.02.011.

[6] J.J. Buccafusco, ed., Methods of Behavior Analysis in Neuroscience, 2nd ed., CRC Press/Taylor & Francis, Boca Raton (FL), 2009. http://www.ncbi.nlm.nih.gov/books/NBK5228/ (accessed June 20, 2020).

[7] A.V. Kalueff, A.M. Stewart, R. Gerlai, Zebrafish as an emerging model for studying complex brain disorders, Trends Pharmacol. Sci. 35 (2014) 63–75. https://doi.org/10.1016/j.tips.2013.12.002.

[8] D. Wahlsten, Mouse Behavior Testing, Academic Press, 2011.

[9] R. Aoki, T. Tsuboi, H. Okamoto, Y-maze avoidance: an automated and rapid associative learning paradigm in zebrafish, Neurosci. Res. 91 (2015) 69–72. https://doi.org/10.1016/j.neures.2014.10.012.

[10] R.M.J. Deacon, J.N.P. Rawlins, T-maze alternation in the rodent, Nat. Protoc. 1 (2006) 7–12. https://doi.org/10.1038/nprot.2006.2.

[11] Y.-H. Kim, K.-S. Lee, Y.-S. Kim, Y.-H. Kim, J.-H. Kim, Effects of hypoxic preconditioning on memory evaluated using the T-maze behavior test, Anim. Cells Syst. 23 (2019) 10–17. https://doi.org/10.1080/19768354.2018.1557743.

[12] Z.K. Varga, Á. Zsigmond, D. Pejtsik, M. Varga, K. Demeter, É. Mikics, J. Haller, M. Aliczki, The swimming plus-maze test: a novel high-throughput model for assessment of anxiety-related behaviour in larval and juvenile zebrafish (Danio rerio), Sci. Rep. 8 (2018) 16590. https://doi.org/10.1038/s41598-018-34989-1.

[13] A.A. Walf, C.A. Frye, The use of the elevated plus maze as an assay of anxiety-related behavior in rodents, Nat. Protoc. 2 (2007) 322–328. https://doi.org/10.1038/nprot.2007.44.

[14] H.A. Bimonte-Nelson, ed., The Maze Book: Theories, Practice, and Protocols for Testing Rodent Cognition, Humana Press, 2015. https://doi.org/10.1007/978-1-4939-2159-1.

[15] S. Kwon, J.-E. Bae, S.-H. Lee, K.-S. Chae, Effects of gravity on positive phototaxis in fruit fly Drosophila melanogaster: Gravity and phototaxis in fruit fly, 46 (2016) 272–277. https://doi.org/10.1111/1748-5967.12173.

[16] R.B. Coleman, K. Aguirre, H.P. Spiegel, C. Pecos, J.A. Carr, B.N. Harris, The plus maze and scototaxis test are not valid behavioral assays for anxiety assessment in the South African clawed frog, J. Comp. Physiol. A Neuroethol. Sens. Neural. Behav. Physiol. 205 (2019) 567–582. https://doi.org/10.1007/s00359-019-01351-3.

[17] T. Lucon-Xiccato, A. Bisazza, Complex maze learning by fish, Anim. Behav. 125 (2017) 69–75. https://doi.org/10.1016/j.anbehav.2016.12.022.

[18] K.A. Connors, T.W. Valenti, K. Lawless, J. Sackerman, E.S. Onaivi, B.W. Brooks, G.G. Gould, Similar anxiolytic effects of agonists targeting serotonin 5-HT1A or cannabinoid CB receptors on zebrafish behavior in novel environments, Aquat. Toxicol. Amst. Neth. 151 (2014) 105–113. https://doi.org/10.1016/j.aquatox.2013.12.005.

[19] J. Sackerman, J.J. Donegan, C.S. Cunningham, N.N. Nguyen, K. Lawless, Long, R.H. Benno, G.G. Gould, Zebrafish Behavior in Novel Environments: Effects of Acute Exposure to Anxiolytic Compounds and Choice of Danio rerio Line, Int. J. Comp. Psychol. 23 (2010) 43–61.

[20] G. de P. Cognato, J.W. Bortolotto, A.R. Blazina, R.R. Christoff, D.R. Lara, M.R. Vianna, C.D. Bonan, Y-Maze memory task in zebrafish (Danio rerio): the role of glutamatergic and cholinergic systems on the acquisition and consolidation periods, Neurobiol. Learn. Mem. 98 (2012) 321–328. https://doi.org/10.1016/j.nlm.2012.09.008.

[21] B.T. Ngoc Hieu, N.T. Ngoc Anh, G. Audira, S. Juniardi, R.A.D. Liman, O.B. Villaflores, Y.-H. Lai, J.-R. Chen, S.-T. Liang, J.-C. Huang, C.-D. Hsiao, Development of a Modified Three-Day T-maze Protocol for Evaluating Learning and Memory Capacity of Adult Zebrafish, Int. J. Mol. Sci. 21 (2020) E1464. https://doi.org/10.3390/ijms21041464.

[22] A. Pilehvar, R.M. Town, R. Blust, The effect of copper on behaviour, memory, and associative learning ability of zebrafish (Danio rerio), Ecotoxicol. Environ. Saf. 188 (2020) 109900. https://doi.org/10.1016/j.ecoenv.2019.109900.

[23] X. Cousin, T. Daouk, S. Péan, L. Lyphout, M.-E. Schwartz, M.-L. Bégout, Electronic individual identification of zebrafish using radio frequency identification (RFID) microtags, J. Exp. Biol. 215 (2012) 2729–2734. https://doi.org/10.1242/jeb.071829.

[24] L. Grossman, A. Stewart, S. Gaikwad, E. Utterback, N. Wu, J. Dileo, K. Frank, P. Hart, H. Howard, A.V. Kalueff, Effects of piracetam on behavior and memory in adult zebrafish, Brain Res. Bull. 85 (2011) 58–63. https://doi.org/10.1016/j.brainresbull.2011.02.008.

[25] C. Vignet, M.-L. Bégout, S. Péan, L. Lyphout, D. Leguay, X. Cousin, Systematic screening of behavioral responses in two zebrafish strains, Zebrafish. 10 (2013) 365–375. https://doi.org/10.1089/zeb.2013.0871.

[26] A. Buatois, S. Nguyen, C. Bailleul, R. Gerlai, Colored-Light Preference in Zebrafish (Danio rerio), Zebrafish. (2021). https://doi.org/10.1089/zeb.2020.1977.

[27] M. Cleal, M.O. Parker, Moderate developmental alcohol exposure reduces repetitive alternation in a zebrafish model of fetal alcohol spectrum disorders, Neurotoxicol. Teratol. 70 (2018) 1–9. https://doi.org/10.1016/j.ntt.2018.09.001.

[28] M.P. Faillace, A. Pisera-Fuster, R. Bernabeu, Evaluation of the rewarding properties of nicotine and caffeine by implementation of a five-choice conditioned place preference task in zebrafish, Prog. Neuropsychopharmacol. Biol. Psychiatry. 84 (2018) 160–172. https://doi.org/10.1016/j.pnpbp.2018.02.001.

[29] S. Zhang, X. Liu, M. Sun, Q. Zhang, T. Li, X. Li, J. Xu, X. Zhao, D. Chen, X. Feng, Reversal of reserpine-induced depression and cognitive disorder in zebrafish by sertraline and Traditional Chinese Medicine (TCM), Behav. Brain Funct. BBF. 14 (2018) 13. https://doi.org/10.1186/s12993-018-0145-8.

[30] R. Benvenutti, M. Marcon, M. Gallas-Lopes, A.J. de Mello, A.P. Herrmann, A. Piato, Swimming in the maze: An overview of maze apparatuses and protocols to assess zebrafish behavior, Neurosci. Biobehav. Rev. 127 (2021) 761–778. https://doi.org/10.1016/j.neubiorev.2021.05.027.

[31] M.B. Orger, G.G. de Polavieja, Zebrafish Behavior: Opportunities and Challenges, Annu. Rev. Neurosci. 40 (2017) 125–147. https://doi.org/10.1146/annurev-neuro-071714-033857.

[32] A. Stewart, S. Gaikwad, E. Kyzar, J. Green, A. Roth, A.V. Kalueff, Modeling anxiety using adult zebrafish: a conceptual review, Neuropharmacology. 62 (2012) 135–143. https://doi.org/10.1016/j.neuropharm.2011.07.037.

[33] R. Gerlai, Learning and memory in zebrafish (Danio rerio), Methods Cell Biol. 134 (2016) 551–586. https://doi.org/10.1016/bs.mcb.2016.02.005.

[34] U.P. Kundap, Y. Kumari, I. Othman, M.F. Shaikh, Zebrafish as a Model for Epilepsy-Induced Cognitive Dysfunction: A Pharmacological, Biochemical and Behavioral Approach, Front. Pharmacol. 8 (2017) 515. https://doi.org/10.3389/fphar.2017.00515.

[35] S. Saleem, R.R. Kannan, Zebrafish: an emerging real-time model system to study Alzheimer’s disease and neurospecific drug discovery, Cell Death Discov. 4 (2018) 45. https://doi.org/10.1038/s41420-018-0109-7.

[36] R. Gerlai, T.L. Poshusta, M. Rampersad, Y. Fernandes, T.M. Greenwood, M.A. Cousin, E.W. Klee, K.J. Clark, Forward Genetic Screening Using Behavioral Tests in Zebrafish: A Proof of Concept Analysis of Mutants, Behav. Genet. 47 (2017) 125–139. https://doi.org/10.1007/s10519-016-9818-y.

[37] G. Bruni, A.J. Rennekamp, A. Velenich, M. McCarroll, L. Gendelev, E. Fertsch, J. Taylor, P. Lakhani, D. Lensen, T. Evron, P.J. Lorello, X.-P. Huang, S. Kolczewski, G. Carey, B.J. Caldarone, E. Prinssen, B.L. Roth, M.J. Keiser, R.T. Peterson, D. Kokel, Zebrafish behavioral profiling identifies multitarget antipsychotic-like compounds, Nat. Chem. Biol. 12 (2016) 559– 566. https://doi.org/10.1038/nchembio.2097.

[38] A. Avdesh, M.T. Martin-Iverson, A. Mondal, M. Chen, S. Askraba, N. Morgan, M. Lardelli, D.M. Groth, G. Verdile, R.N. Martins, Evaluation of color preference in zebrafish for learning and memory, J. Alzheimers Dis. JAD. 28 (2012) 459–469. https://doi.org/10.3233/JAD-2011-110704.

[39] Y.H. Kim, K.S. Lee, A.R. Park, T.J. Min, Adding preferred color to a conventional reward method improves the memory of zebrafish in the T-maze behavior model, Anim. Cells Syst. 21 (2017) 374–381. https://doi.org/10.1080/19768354.2017.1383938.

[40] E.A. Osipova, V.V. Pavlova, V.A. Nepomnyashchikh, V.V. Krylov, Influence of magnetic field on zebrafish activity and orientation in a plus maze, Behav. Processes. 122 (2016) 80–86. https://doi.org/10.1016/j.beproc.2015.11.009.

[41] J.-S. Park, J.-H. Ryu, T.-I. Choi, Y.-K. Bae, S. Lee, H.J. Kang, C.-H. Kim, Innate Color Preference of Zebrafish and Its Use in Behavioral Analyses, Mol. Cells. 39 (2016) 750–755. https://doi.org/10.14348/molcells.2016.0173.

[42] R. Gerlai, Reproducibility and replicability in zebrafish behavioral neuroscience research, Pharmacol. Biochem. Behav. 178 (2019) 30–38. https://doi.org/10.1016/j.pbb.2018.02.005.

[43] J. Oliveira, M. Silveira, D. Chacon, A. Luchiari, The zebrafish world of colors and shapes: preference and discrimination, Zebrafish. 12 (2015) 166–173. https://doi.org/10.1089/zeb.2014.1019.

[44] C. Lieggi, A.V. Kalueff, C. Lawrence, C. Collymore, The Influence of Behavioral, Social, and Environmental Factors on Reproducibility and Replicability in Aquatic Animal Models, ILAR J. (2020). https://doi.org/10.1093/ilar/ilz019.

[45] M. Sison, R. Gerlai, Associative learning in zebrafish (Danio rerio) in the plus maze, Behav. Brain Res. 207 (2010) 99. https://doi.org/10.1016/j.bbr.2009.09.043.

[46] A. Facciol, S. Tran, R. Gerlai, Re-examining the factors affecting choice in the light-dark preference test in zebrafish, Behav. Brain Res. 327 (2017) 21– 28. https://doi.org/10.1016/j.bbr.2017.03.040.

[47] C. Maximino, T. Marques de Brito, C.A.G. de M. Dias, A. Gouveia, S. Morato, Scototaxis as anxiety-like behavior in fish, Nat. Protoc. 5 (2010) 209–216. https://doi.org/10.1038/nprot.2009.225.

[48] E.L. Serra, C.C. Medalha, R. Mattioli, Natural preference of zebrafish (Danio rerio) for a dark environment, Braz. J. Med. Biol. Res. Rev. Bras. Pesqui. Medicas E Biol. 32 (1999) 1551–1553. https://doi.org/10.1590/s0100-879X1999001200016.

[49] S.J. Dahlbom, T. Backström, K. Lundstedt-Enkel, S. Winberg, Aggression and monoamines: effects of sex and social rank in zebrafish (Danio rerio), Behav. Brain Res. 228 (2012) 333–338. https://doi.org/10.1016/j.bbr.2011.12.011.

[50] C.L. Rambo, R. Mocelin, M. Marcon, D. Villanova, G. Koakoski, M.S. de Abreu, T.A. Oliveira, L.J.G. Barcellos, A.L. Piato, C.D. Bonan, Gender differences in aggression and cortisol levels in zebrafish subjected to unpredictable chronic stress, Physiol. Behav. 171 (2017) 50–54. https://doi.org/10.1016/j.physbeh.2016.12.032.

[51] R. Genario, M.S. de Abreu, A.C.V.V. Giacomini, K.A. Demin, A.V. Kalueff, Sex differences in behavior and neuropharmacology of zebrafish, Eur. J. Neurosci. 52 (2020) 2586–2603. https://doi.org/10.1111/ejn.14438.

[52] N. Percie du Sert, V. Hurst, A. Ahluwalia, S. Alam, M.T. Avey, M. Baker, W.J. Browne, A. Clark, I.C. Cuthill, U. Dirnagl, M. Emerson, P. Garner, S.T. Holgate, D.W. Howells, N.A. Karp, S.E. Lazic, K. Lidster, C.J. MacCallum, M. Macleod, E.J. Pearl, O.H. Petersen, F. Rawle, P. Reynolds, K. Rooney, E.S. Sena, S.D. Silberberg, T. Steckler, H. Würbel, The ARRIVE guidelines 2.0: Updated guidelines for reporting animal research, BMC Vet. Res. 16 (2020). https://doi.org/10.1186/s12917-020-02451-y.

[53] American Veterinary Medical Association, AVMA Guidelines for the Euthanasia of Animals: 2020 Edition, (2020). https://www.avma.org/sites/default/files/2020-01/2020-Euthanasia-Final-1-17-20.pdf.

[54] Z.A. Bault, S.M. Peterson, J.L. Freeman, Directional and color preference in adult zebrafish: Implications in behavioral and learning assays in neurotoxicology studies, J. Appl. Toxicol. JAT. 35 (2015) 1502–1510. https://doi.org/10.1002/jat.3169.

[55] E. Choleris, L.A.M. Galea, F. Sohrabji, K.M. Frick, Sex differences in the brain: Implications for behavioral and biomedical research, Neurosci. Biobehav. Rev. 85 (2018) 126–145. https://doi.org/10.1016/j.neubiorev.2017.07.005.

[56] J. Oliveira, M. Silveira, D. Chacon, A. Luchiari, The zebrafish world of colors and shapes: preference and discrimination, Zebrafish. 12 (2015) 166–173. https://doi.org/10.1089/zeb.2014.1019.

[57] A. Buatois, S. Nguyen, C. Bailleul, R. Gerlai, Colored-Light Preference in Zebrafish (Danio rerio), Zebrafish. 18 (2021) 243–251. https://doi.org/10.1089/zeb.2020.1977.

[58] E. Angiulli, V. Pagliara, C. Cioni, F. Frabetti, F. Pizzetti, E. Alleva, M. Toni, Increase in environmental temperature affects exploratory behaviour, anxiety and social preference in Danio rerio, Sci. Rep. 10 (2020). https://doi.org/10.1038/s41598-020-62331-1.

[59] R.E. Blaser, D.B. Rosemberg, Measures of Anxiety in Zebrafish (Danio rerio): Dissociation of Black/White Preference and Novel Tank Test, PLoS ONE. 7 (2012). https://doi.org/10.1371/journal.pone.0036931.

[60] C. Maximino, B. Puty, K.R. Matos Oliveira, A.M. Herculano, Behavioral and neurochemical changes in the zebrafish leopard strain, Genes Brain Behav. 12 (2013) 576–582. https://doi.org/10.1111/gbb.12047.

[61] R.E. Blaser, L. Chadwick, G.C. McGinnis, Behavioral measures of anxiety in zebrafish (Danio rerio), Behav. Brain Res. 208 (2010) 56–62. https://doi.org/10.1016/j.bbr.2009.11.009.

[62] L.D.P. Magno, A. Fontes, B.M.N. Gonçalves, A. Gouveia, Pharmacological study of the light/dark preference test in zebrafish (Danio rerio): Waterborne administration, Pharmacol. Biochem. Behav. 135 (2015) 169–176. https://doi.org/10.1016/j.pbb.2015.05.014.

[63] P.J. Steenbergen, M.K. Richardson, D.L. Champagne, The use of the zebrafish model in stress research, Prog. Neuropsychopharmacol. Biol. Psychiatry. 35 (2011) 1432–1451. https://doi.org/10.1016/j.pnpbp.2010.10.010.

[64] S.D. Córdova, T.G. Dos Santos, D.L. de Oliveira, Water column depth and light intensity modulate the zebrafish preference response in the black/white test, Neurosci. Lett. 619 (2016) 131–136. https://doi.org/10.1016/j.neulet.2016.03.008.

[65] A. Facciol, M. Iqbal, A. Eada, S. Tran, R. Gerlai, The light-dark task in zebrafish confuses two distinct factors: Interaction between background shade and illumination level preference, Pharmacol. Biochem. Behav. 179 (2019) 9–21. https://doi.org/10.1016/j.pbb.2019.01.006.

[66] J.F. Stephenson, K.E. Whitlock, J.C. Partridge, Zebrafish preference for light or dark is dependent on ambient light levels and olfactory stimulation, Zebrafish. 8 (2011) 17–22. https://doi.org/10.1089/zeb.2010.0671.

